# Tuberculosis at farmer-cattle interface in the rural villages of South Gondar Zone of northwest Ethiopia

**DOI:** 10.1101/605154

**Authors:** Amir Alelign, Aboma Zewude, Beyene Petros, Gobena Ameni

## Abstract

**Background:** Tuberculosis (TB) has been an important public health concern in Ethiopia, particularly at areas of human-animal intersection. However, limited epidemiological information is available in this respect in the country. Therefore, the present study was conducted to investigate the transmission of TB at human-cattle interface, associated risk factors and public awareness about the disease at South Gondar Zone, northwest Ethiopia.

**Methods:** A cross sectional study was conducted between March 2015 and April 2018 on 186 farmers and 476 cattle in South Gondar Zone, northwest Ethiopia. Bacteriological examination, region of difference (RD) 9 based polymerase chain reaction (PCR), single intradermal comparative tuberculin test (SIDCTT) and questionnaire were used for undertaking this study.

**Results:** Culture positivity in farmers was 59.7% (111/186) and all the culture positive isolates were *M. tuberculosis*. About 68% (74/111) of culture positive respondents did not know about the transmission of TB from cattle to human or vice versa. The animal and herd prevalence of bovine TB were 1.5% (7/476) and 7.4% (7/95) respectively. The odd of bovine TB in cattle owned by TB positive households was slightly higher than those owned by TB free households (adjusted odds ratio, AOR=1.39; 95% CI: 0.31-7.10; p = 0.76).

**Conclusion:** Although SIDCTT reactivity was slightly higher in cattle owned by TB positive households, all the human isolates were *M. tuberculosis* and no *M. bovis* was isolated from farmers, which could be due to the low prevalence of bovine TB in the area.

## Introduction

Tuberculosis (TB) is an infectious disease of humans and animals. The most common cause of human TB is known to be *Mycobacterium tuberculosis* (*M. tuberculosis*), and the main cause of TB in animals is *Mycobacterium bovis* (*M. bovis*). Zoonotic TB is a form of TB that is caused by *M. bovis* and transmitted from animals to humans. While reverse zoonotic TB is the form of TB that is caused by *M. tuberculosis* and is transmitted from humans to animals. *M. bovis* often causes extra pulmonary and can also cause pulmonary TB which is clinically indistinguishable from TB caused by *M. tuberculosis*. In 2016, an estimated 147, 000 new cases of zoonotic TB were reported globally, and 12, 500 deaths due zoonotic TB. The highest burden of zoonotic TB was reported from the African, followed by the South-East Asian region [1].

The transmission of TB from cattle to human is due to consumption of raw/undercooked infected animal products such as milk and meat or through inhalation due to close contact bet -ween the cattle and humans. It is estimated that in countries where pasteurization of milk is rare and bovine TB is common, 10 to 15% human cases of TB are caused by *M. bovis* [2]. According to some studies from Tanzania, Nigeria and Uganda, *M. bovis* accounted for 20% or more of the MTBC isolated from human TB cases [3-5].

Data on TB transmission at the human-animal interface are important in designing a ‘one health’ approach for the control of the disease, particularly, rural settings. In the Amhara Region, north-western Ethiopia, information on the public health risk of zoonotic TB is scarce. Few cross sectional studies conducted in cattle reported animal prevalence of 3.55% and 8.7% [7, 8]. In addition, *M. bovis* was isolated from humans in north-western Ethiopia [8, 9].

In South Gondar Zone, Amhara Region, north-western Ethiopia, the practices of inhabitants could promote the transmission of TB from cattle to humans or vice versa. However, there is scarcity of epidemiological data on public awareness, risk factors and transmission of TB between humans and their cattle in the Zone. Hence, the present study was conducted to investigate the public awareness, risk factors to BTB and the potential transmission of TB from cattle to humans or vice versa in South Gondar Zone of north-western Ethiopia.

## Materials and methods

### Study area

The study was conducted in South Gondar Zone, north-western Ethiopia. The Zone is located in the Amhara Region, 660 km north east of Addis Ababa, the Capital of Ethiopia. The Zone is known for its diverse topography ranging from flat and low grazing land to high cold mountains. The altitude of the Zone ranges from1500 to 3,600 meters above sea level. The average yearly rainfall of the Zone ranged from 700-mm to 1300 mm in while the average daily temperature was 17° C in 2017. South Gondar Zone consisted of 10 districts and covers a total area of 14 320 square km. According to the Central Statistical Agency of Ethiopia, the Zone has a total population of 2,051,738 and a population density of 145.56 [10]. Majority of the population (90.47%) of the Zone were rural inhabitants. A total of 468,238 households were counted in this Zone, which resulted in an average of 4.38 persons to a household [10].

The majority of the population has depended on subsistence farming and dairy cattle rearing [11]. The Zone has been known for its indigenous milk cattle such as the ‘Fogera’ and ‘Dera’ cattle. Dairying is commonly practiced using small herd size.

### Study population and sampling

Human TB cases were recruited from districts health centres, peripheral health posts, Zonal and regional government as well as private referal hospitals (Debretabor, Felegehiwot and Gambi hosptals). The human cases included those active TB patients who visited the health facilities seeking for TB-treatment. The control human subjects were households who did not have TB cases for the last five years and residing in the same villages with the households which have TB cases in their family members. Cattle owned by both TB positive households and TB negative households were tested for bovine TB.

### General inclusion and exclusion criteria

Small scale dairy farmers who were permanently living in the zone for at least two months prior to the study, aged five years and above at the time of the study as well as who were consented and willing to participate in the study were included. Whereas, individuals who had no association with cattle, those who were not willing to participate, aged below five years and seriously ill ones were excluded from participating in the study. Both local and cross breed cattle older than six months and owned by households with active TB cases and TB free households were included. Young cattle with age less than six months, clinically sick ones, and cows one month pre-and post partum were excluded from the study.

### Study design and sample size determination

A comparative cross sectional study was conducted between December 2015 to February 2018 The sample size for this study was calculated with the assumption of a 15% bovine TB prevalence in TB positive households and a 5% bovine TB prevalence among TB free households, a 95% confidence interval (CI) with a power of 80%, a ratio of cases to comparison groups of 3:1 (based on findings of 15.3% bovine TB among TB positive households and a 5.9% BTB among TB free households [8]. Adding a 10% non-response rate, the required sample size was 105 for TB positive households and 35 TB for free households. However, in the present study, due to several limitations we have only managed to reach out only 63 TB positive and 32 TB negative households. Based on this, a total of 476 cattle (315 from TB positive households and 161 from TB free households) were tested for bovine TB.

## Data collection

### Questioner survey

All human study participants were interviewed with questionnaire in order to assess their awareness and practices about the zoonotic transmission of TB, and the possible risk factors for the disease. Their socio-demographic characteristics, knowledge, attitudes and practices towards TB including their consumption habits of raw milk/meat as well as husbandry practices were asked. Moreover, all herd owners of tuberculin tested cattle were interviewed for possible risk factors of TB positive cattle. Data on age and sex of individual animal, herd size, origin, breed type, recent introduction of new animals into the herd, and keeping of different livestock together were collected.

### Laboratory methods

After informed consent was obtained, two sputum samples (on-spot and morning) from suspected pulmonary TB (PTB) patients and fine needle aspirate (FNA) samples from suspect -ed extrapulmonary TB (EPTB) patients were collected by trained laboratory technicians and pathologists, respectively.

The samples were first digested and concentrated/decontaminated by the N-acetyl-L-cysteine-Sodium hydroxide (NALC-NaOH) method. Smears of the final deposits from the various specimens were stained by the Ziehl-Neelsen (ZN) method and examined under oil immersion using a binocular light microscope. All smear positive TB samples were stored at +4°C at the study site and then transported to Regional Health Research Llaboratory Center (Bahirdar) and kept at +4 °C until bacteriological culture was performed.

Similarly, FNA specimens were collected by pathologists and stored in cryo-tubes in phosphate buffer saline (PBS) with pH 7.2. ZN staining was performed. AFB-positive specimens were stored at −20 °C until mycobacterial culture was performed.

The samples were processed for culturing according to the standard methods described earlier [12, 13]. Both sputum and FNA samples were cultured at the Bahir Dar Regional Health Research Laboratory Centre.

To differentiate *M. tuberculosis* from other members of the *M. tuberculosis* complex (MTBC) species, RD9 based PCR was performed according to protocols previously described [14]. *M. tuberculosis* H37Rv, *M. bovis* bacille Calmette-Guérin (BCG) were included as positive controls and water was used as a negative control. Interpretation of the result was based on bands of different sizes (396 base pairs (bp) for *M. tuberculosis* and 375bp for *M. bovis*) as previously described [15].

### Animal study

#### Single intradermal comparative tuberculin test

Single intradermal comparative tuberculin test (SIDCTT) was performed for detecting TB in cattle. SIDCTT was done using both bovine and avian purified protein derivatives (PPDs) (Prionics Lelystad B. V., The Netherlands). The tuberculin test measures the hypersensitivity reaction on the skin due to the administered antigens (PPDs).

Two sites were shaved in the middle of the side of the neck, one above the other, separated by about 12 cm for injection of the two PPDs. The thickness of the skin fold at both injection sites were measured using a calliper and recorded before injection. The sites were injected with 0.1ml (2000IU) aliquot of bovine PPD and 0.1ml (2500IU) of avian PPD interdermally. After 72 hours the two sites were measured for change in skin thickness, and the result was interpreted according international [16] and local [17] criteria.

## Data Analysis

Data were entered in to Excele file format and transferred in to SPSS software version 25 for statistical analysis. Descriptive statistics were used to depict the demographic variables. Chi-square (χ^2^) test was used to test differences in proportions and the association between categorical variables with raw milk consumption habit. Bivariate and multivariate logistic regressions were used to determine the association between background variables with awareness of zoonotic TB and tuberculin reactivity in cattle. Results were considered statistically significant whenever p-value was less than 5%.

## Ethics approval and consent to participate

Ethical clearance for the study was obtained from the Ethical Committee of Addis Ababa University, Department of Microbial, Cellular and Molecular Biology (Ref. CNSDO/491/07/ 15). In addition, written permission was sought from the Amhara Regional Health Bureau Research Ethical Committee (Ref. HRTT/1/271/07). Each study participant was consented with a written form and agreed to participate in the study after a clear explanation of the study objectives and patient data confidentiality (S1 Appendix). In case of participants under the age of 18 years, consent was obtained from their parents/guardians. The animals used in this study were privately owned by the study participants and a written consent was sought from the owners to take samples from the animals. A high standard veterinary care was taken to minimize cattle suffering during tuberculin test.

## Results

### Tuberculosis in farmers

Culture positivity was obtained in 59.7% (111/186) of the active TB cases. Of which, 59.5% (66/111) was isolated from extra pulmonary TB (EPTB) patients. The molecular typing of culture positive isolates using RD9-based PCR revealed that all isolates had intact RD9 locus and were subsequently classified as *M. tuberculosis.* No *M. bovis* was detected (Fig 1).

**Fig 1.**
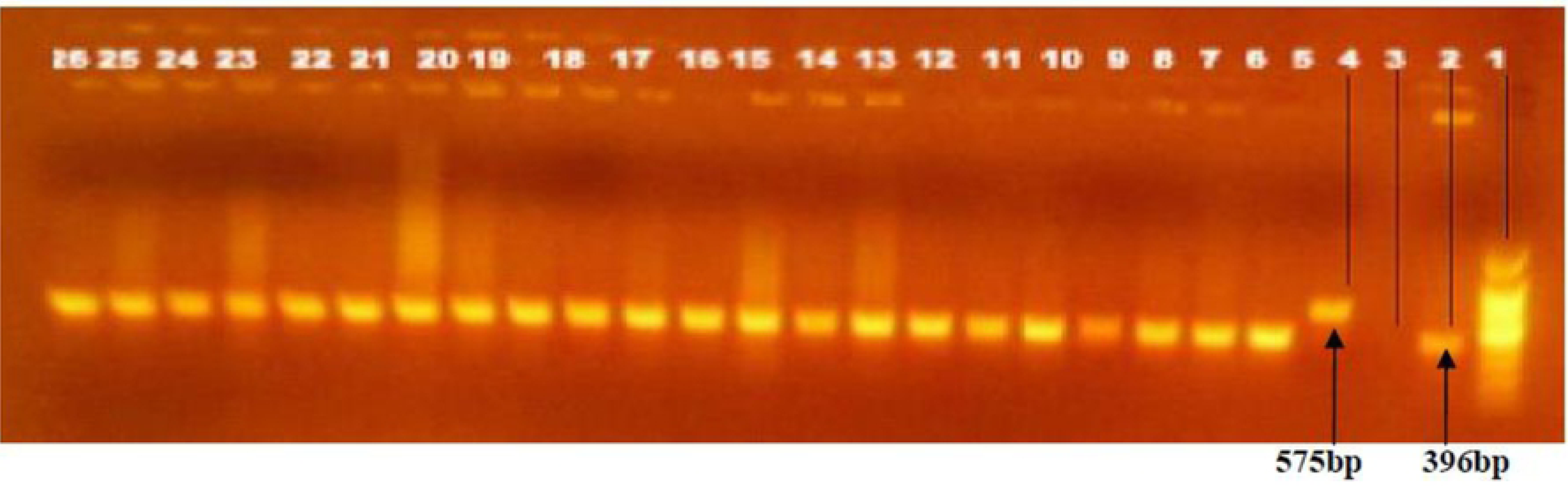
Electrophoretic separation of PCR products by RD9 deletion typing. The figure repre -sents only for 22 mycobacteria isolates from sputum and FNA human samples: Lane 1 DNA ladder, 2 *M. tuberculosis* control (396 base pair), 3 molecular grade water (negative control), 4 *M. bovis* control (575 bp), lane 5 26 are culture isolates of *M. tuberculosis* from human tuberculosis patients designated with their sample code as: 5FE1, 6FE2,7FE3,8FE4, 9FE7, 10FE11, 11FE12,12FE15, 13FE16,14FE19,15FE21, 16FE23, 17FE24, 18FE25, 19FE27, 20FE28, 21FE31, 22FE33, 23FE36, 24FE38, 25FE47, 26FE48.

#### Awarness on zoonotic transmission of TB and food consumption habit

About 68% (74/111) of the respondents did not know about the transmission of TB from cattle to human or vice versa. About 69% (77/111) of the respondents had habit of consuming raw milk and other uncooked dairy products (Table 1).

**Table 1.** Association of demographic factors with awarness to zoonotic transmission of TB among AFB culture positive TB patients (N=111), South Gondar Zone, northwest Ethiopia (2015-2017).

| Demographic factors | Number of respondents (%) | Aware of zoonotic TB |  | COR (95% CI) | AOR (95% CI) | p-value |
| --- | --- | --- | --- | --- | --- | --- |
|  |  | Yes ((%) | No (%) |  |  |  |
| Patient origin |  |  |  |  |  |  |
| Dera | 26 (23.4) | 6 (23.1) | 20 (76.9) | 1.00 | 1.00 |  |
| Ebinat | 10 (9) | 2 (20) | 8 (80) | 0.83(0.13-5.03) | 0.66 (0.39-1.28) | 0.842 |
| Este | 15 (13.5) | 1(6.7) | 14 (93.3) | 0.23(0.02-2.20) | 0.25 (0.01-2.53) | 0.178 |
| Farta | 9 (8.1) | 2 (22.2) | 7(77.8) | 0.95(0.15-5.86) | 0.63 (0.02-5.53) | 1.00 |
| Fogera | 35 (31.5) | 13(37.1) | 22(62.9) | 1.96(0.62-6.16) | 1.99 (0.58-8.02) | 0.240 |
| Gayint | 6 (5.4) | 3(50) | 3(50) | 3.33(0.52-21.03) | 3.24 (0.31-21.0) | 0.186 |
| Libo kemkem | 7(6.3) | 5(71.4) | 2 (28.6) | 11.66(1.89-71.79) | 11.84 (1.67-73.2) | 0.003 |
| Simada | 3(2.7) | 2 (66.7) | 1(39.3) | 10.0(0.87-114.74) | 10.47(0.72-116.3) | 0.038 |
| Age group (year) |  |  |  |  |  |  |
| <18 | 10(9) | 5(50) | 5(50) | 1.00 | 1.00 |  |
| 18-30 | 42(37.8) | 17(40.5) | 25(59.5) | 0.68(0.17-2.71) | 0.64 (0.13-2.94) | 0.583 |
| 31-43 | 35(31.5) | 8(22.9) | 27(77.1) | 0.29(0.06-1.28) | 0.32 (0.04-1.42) | 0.094 |
| 44-56 | 18(16.2) | 5(27.8) | 13(72.2) | 0.38(0.07-1.92) | 0.46 (0.09-2.31) | 0.239 |
| >56 | 6(5.4) | 2(33.3) | 4(66.7) | 0.50(0.06-4.09) | 0.51 (0.07-5.21) | 0.515 |
| Sex |  |  |  |  |  |  |
| Male | 58(52.3) | 18(31.0) | 40(69.0) | 1.00 | 1.00 |  |
| Female | 53(47.7) | 19(35.8) | 34(64.2) | 1.24(0.56-2.73) | 1.16 (0.49-2.97) | 0.590 |
| Education status |  |  |  |  |  |  |
| Illiterate | 73(65.8) | 22(30.1) | 51(69.9) | 1.00 | 1.00 |  |
| Adult education | 5(4.5) | 1(0.2) | 4(99.8) | 0.57(0.06-5.48) | 0.23 (0.04-5.53) | 0.630 |
| Primary level | 18(16.2) | 4(22.2) | 14(77.8) | 0.66(0.19-2.24) | 0.62 (0.08-2.62) | 0.505 |
| Secondary level | 11(9.9) | 7(63.6) | 4(36.4) | 4.05(1.07-15.28) | 4.16 (1.05-15.57) | 0.029 |
| Higher level | 4(3.6) | 3(0.75) | 1(0.25) | 6.95(0.68-70.60) | 6.52 (0.62-70.71) | 0.062 |
| Raw milk consumption |  |  |  |  |  |  |
| Yes | 77(69.4) | 27(35.1) | 50(64.9) | 1.00 | 1.00 |  |
| No | 34(30.6) | 10(29.4) | 26 (70.6) | 0.71(0.29-1.69) | 0.71 (0.26-1.88) | 0.441 |
| Patient category |  |  |  |  |  |  |
| New | 91(82.0) | 29(31.9) | 62(68.1) | 1.00 | 1.00 |  |
| Retreatment | 20(18.0) | 8(40.0) | 12(60.0) | 1.42(0.52-3.86) | 1.38 (0.42-3.93) | 0.484 |

COR: crude odds ratio, AOR: adjusted odds ratio, CI: confidence interval.

The logistic regression, taking log-odds of awarness about zoonotic transmission as an outcome variable, resulted patient origin and educational status were observed to be significantly associated (p<0.05) (Table 1). The study participants in Libo kemkem and Simada were 11 and 10 times more aware about the zoonotic tramnsmission of TB as compared to those of Dera District (Libo kemkem vs. Dera AOR= 11.84; 95% CI: 1.67-73.2; p = 0.003) and (Simada vs. Dera AOR= 10.47; 95% CI: 0.72-116.3; p = 0.038). The odds of having awareness on zoonotic transmission of TB was higher among individuals with secondary school educational level (AOR=4.16; 95% CI: 1.05-15.57; p = 0.029) compared to those of illiterates (Table 1).

However, other patient characteristics such as age groups, sex, TB history in the family, raw milk consumption habit, and patient category (new or retreatment cases) were not significantly associated with particpants’ over all awarness about zoonotic transmission of TB (Table 1).

### Tuberculosis in cattle

#### Characteristics of the study cattle

The majority of the cattle were females accounting for 54.2% (258/476) of the study cattle. Cattle within the age range of 5-10 years had the greatest share (47%) from the total cattle tested with a mean age of 5.5 years. Many of the cattle were Zebu breed while only 6.1% (29/476) of them were cross breed (Table 2).

**Table 2.** Association of host risk factors with bovine tuberculin test reactivity in cattle based on a >2mm cutoff value, South Gondar Zone, northwest Ethiopia (2015-2018).

| Characteristics | BTB (n=476) |  | Total (%) | COR (95% CI) | AOR (95% CI) | p-value |
| --- | --- | --- | --- | --- | --- | --- |
|  | Positive | Negative |  |  |  |  |
| **Age of cattle (years)** |  |  |  |  |  |  |
| <5 | 1 | 167 | 215(45.2) | 1 |  |  |
| 5-10 | 5 | 203 | 224 (47.0) | 4.11 (0.47- 35.55) | 3.10 (0.35-35.69) | 0.16 |
| >10 | 1 | 106 | 37 (7.8) | 1.58 (0.09-25.45) | 1.69 (0.16-25.93) | 0.74 |
| **Sex** |  |  |  |  |  |  |
| Male | 2 | 215 | 217 (45.6) | 1 |  |  |
| Female | 5 | 254 | 259 (54.4) | 2.11(0.40-11.01) | 2.16 (0.35-12.21) | 0.36 |
| **Breed type** |  |  |  |  |  |  |
| Local | 6 | 441 | 447 (94.0) | 1 |  |  |
| Cross | 1 | 28 | 29 (6.0) | 2.62 (0.30-22.56) | 2.67 (0.25-23.74) | 0.36 |
| **Source** |  |  |  |  |  |  |
| Homebred | 5 | 364 | 368 (77.3) | 1 |  |  |
| Purchased | 2 | 105 | 108 (22.7) | 1.38 (0.27-7.25) | 1.38 (0.17-7.55) | 0.69 |
| **Body condition** |  |  |  |  |  |  |
| Poor | 1 | 196 | 197 (41.4) | 1 |  |  |
| Medium | 3 | 204 | 207(43.5) | 2.88 (0.29-27.94) | 3.00 (0.27-28.38) | 0.33 |
| Good | 3 | 69 | 72 (15.1) | 8.52 (0.87-83.29) | 8.53 (0.85-83.34) | 0.02 |
| **Household TB status** |  |  |  |  |  |  |
| Negative | 2 | 159 | 161(33.8) | 1 |  |  |
| Positive | 5 | 310 | 315(66.2) | 1.28 (0.24-6.68) | 1.39 (0.31-7.10) | 0.76 |

BTB: bovine tuberculosis, n: number of total cattle tested. COR: crude odds ratio, AOR: Adjusted odds ratio, CI: confidence interval.

#### Prevalence of TB in cattle

Animal prevalence was 1.6% (5/315) and 1.2% (2/161) at ≥2mm cut-off value in TB positive and TB free households, respectively. Using the same cut-off value, 7.9% (5/63) and 6.3% (2/32) herd prevalence was recorded in cattle owned by TB positive and TB free households, respectively. The overall animal and herd prevalence were 1.5% (7/476 and 7.4% (7/95), respectively. However, non of the tested cattle were positive for bovine TB at the international cut-off value of >4mm.

### Risk factors for bovine TB

Risk factor analysis to the occurrence of bovine TB in cattle revealed that age groups between 5-10 years were more reactive to tuberculin test than younger age groups (AOR=3.1; 95% CI: 0.35-35.69; p=0.16). Cattle with apparently good (AOR= 8.53; 95% CI: 0.85-83.34; p =0.02) and medium (AOR= 3.00; 95% CI: 0.27-28.38; p = 0.33) body conditions were more likely to be reactive to the tuberculin test as compared to those with apparently poor body condition, and the difference was statistically significant (p<0.05) (Table 2). Although the difference was not statistically significant (p>0.05, the odd of bovine TB that cattle owned by TB positive cases was slightly higher than those owned by TB free households (AOR=1.39; 95% CI: 0.31-7.10; p = 0.76). Despite the observed differences, sex, breed type, source of cattle and households TB status were not significantly associated with the occurrence of BTB in the present study (Table 2).

### Zoonotic transmission of TB

In the present study, molecular typing of culture positive isolates using RD9-based PCR confirmed that all the human isolates were *M. tuberculosis* (Fig 1). Furthermore, no *M. bovis* was detected even from those TB patients who owned tuberculin reactor cattle. Hence, this study did not revealed evidence of direct transmission of tuberculosis from cattle to their closely associated owners.

## Discussion

The identification of *M. tuberculosis* as the only *Mycobacterium* species in the present study, using RD9-based PCR, was in agreement with previous reports in other parts of Ethiopia in which all or the majority of the isolates found from human TB cases were *M. tuberculosis* [7, 18-20]. In contrast, previous studies conducted in large scale commercial farms and pastoral communities in lowland areas suggested the contribution of *M. bovis* to the overall burden of TB in humans [21, 22]. The reason for the difference in *Mycobacterium* species prevalence in this study and previous studies might be due to the low TB infection rate in cattle owned by smallholder farmers that participated in the present study.

Although zoonotic transmission of *M. bovis* from cattle to famers was expected, all the human isolates were *M. tuberculosis*. Nonetheless, previous study conducted in and around Bahir Dar City [9], Borena Zone [21] and Afar Region reported the isolation of *M. bovis* from human TB cases. It has been well established in the literature that the prevalence of human TB caused by *M. bovis* in specific geographic region is directly proportional to the prevalence of bovine TB in that specific geographic region [23]. In the present study, the prevalence of bovine TB was very low and hence the chance of its transmission to humans is minimal.

Nevertheless, the awareness of farmers about zoonotic transmission of TB was low and thus was similar with the magnitude of awareness recorded by previous studies conducted in Ethiopia [9, 22, 24] and in others including Zambia and Zimbabwe [25, 26]. The poor awarness of farmers on the transmission of zoonotic TB to them could pose risk of infection by zoonotic pathogens including *M. bovis.* On the other hand, in contrast to the low awareness of the farmers included in the present study, farmers in Cameroon and Malawi had good awareness on the zoonotic TB and its transmission [27, 28].

Although *M. bovis* was not isolated from the farmers with active TB, majority of culture posistive TB patients had the habit of consuming raw milk. This observation is similar with the observation of previous studies conducted in different parts of Ethiopia [9, 29-31]. The higher preference of raw milk consumption in Ethiopia could be associated to culture, its taste, availability in the local market, an easy access from a door to door supply by farmers and lower price [32].

The animal and herd prevalence of bovine TB at a severe cut-off value of SIDCTT were low in South Gondar Zone of north-western Ethiopia. In agreement to the prevalence report of this study, low prevalence of bovine TB was reported in and around Bahir Dar City and Yeki District of southern Ethiopia [7, 33]. On the other hand, higher prevalence of bovine TB was reported in and around other cities of Ethiopia [17, 34-37]. These variations in the prevalence of bovine TB are associated with the breed of cattle kept and the type of husbandry under which the cattle are kept. Previous studies in Ethiopia have indicated that *Bos taraus* breed is more susceptible to bovine TB as compared to *Bos indicus* breed [38]. Moreover, it was observed that cattle kept in intensive farms are more susceptible to bovine TB as compared to cattle kept in extensive farms [38]. In addition, it was well established that the prevalence of bovine TB is directly associated with the herd size. Thus, the observation of low prevalence bovine TB in the present study is not surprising as all the study cattle were *Bos indicus* and were also kept in extensive farming; both of which do not favour the occurrence and transmission of bovine TB. Furthermore, all the herds included in the present study were small and thus did not favour the transmission of bovine TB.

Nevertheless, although the overall prevalence of bovine TB recorded by the present study was low, it was relatively higher in cattle owned by TB positive households than cattle owned by TB free households. But the difference of prevalence between the two groups was not statistically significant. Other earlier studies reported higher prevalence of bovine TB in cattle owned by households with active TB cases than TB free households [8, 35, 39, 40]. Such observation could suggest the existence of TB transmission between cattle and their owners. The transmission could be zoonotic (transmission of *M. bovis* from cattle to humans) or reverse zoonotic (transmission of *M. tuberculosis* from humans to cattle). In the present study, isolation of mycobacteria was not done from cattle and diagnosis of bovine TB was made by SIDCTT. On the other hand, all the human isolates were *M. tuberculosis*. This could imply that *M. tuberculosis* might have been transmitted to cattle from their owners and positivity to SIDCTT was due to sensitization to infection with *M. tuberculosis* as it was observed earlier by other authors (14, 39, 41).

The present study has some limitation in conducting pathological examination and there by strain identification of mycobacteria isolates from tuberculin reactor cattle.

## Conclusion

All the human isolates recovered from farmers with active TB cases were *M. tuberculosis* and no *M. bovis* was isolated from farmers. Moreover, the overall prevalence of bovine TB in the area was low; but it was slightly higher in cattle owned by households with active TB cases than in cattle owned with active TB free households; which could suggest the presence of zoonotic and or reverse zoonotic transmission of TB between cattle and their owners. This could also be exacerbated by the low level of awareness of the farmers on the transmission of mycobacterial species between cattle and their owners.

## Acknowledgements

We also would like to thank all laboratory working staff and administrators at the Regional Health research Center, Bahirdar and ALIPB, Addis Ababa University, for their support in the development of this study. We also greatly thank health officials, laboratory workers and study participants in the study area, without whom this study would have not been completed.

## Supporting information

**S1 Appendix: Informed consent form for the study participants.** The form consists of detailed information about the purpose of the study and a written consent form for the study participants who attend health facilities during the study period in South Gondar Zone, northwest Ethiopia.

